# Effects of aging on multiple object tracking under normal and altered viewing conditions

**DOI:** 10.64898/2026.06.25.734471

**Authors:** Célia Michaud, Robin Baurès, Vincent Soler, Yves Trotter, Victor Vattier, Maxime Rosito, Carole Peyrin, Benoit R. Cottereau

**Affiliations:** Univ. Toulouse, CNRS, CerCo, Toulouse, France; Univ Paul Valery of Montpellier, EPSYLON EA, 4556 Montpellier, France; EuroMov Digital Health in Motion, Univ Montpellier, IMT Mines Ales, Montpellier, France; Univ. Grenoble Alpes, Univ. Savoie Mont Blanc, CNRS, LPNC, 38000 Grenoble, France; IPAL, CNRS IRL 2955, Singapore

**Keywords:** Motion perception, Aging, Multiple object tracking, Eye movements, Gaze-contingent paradigm, Simulated visual field loss

## Abstract

Multiple object tracking (MOT) is a core function of dynamic visual attention that relies on the ability to simultaneously monitor several moving objects. Although MOT performance is known to decline with age, and to depend on efficient oculomotor strategies, how these processes interact across the adult lifespan and under degraded visual input remains poorly understood. Here, we examined the effects of aging on MOT under normal and gaze-contingent viewing conditions simulating central and peripheral visual field loss. Sixty participants aged 20-80 years completed a MOT task while eye movements were recorded, enabling characterization of performance and oculomotor behavior across five viewing conditions. Behavioral results revealed a continuous decline in tracking performance across adulthood, indicating a graded rather than categorical effect of age. Performance was strongly reduced by visual-field restrictions, with the largest impairments under central vision occlusion. Eye-tracking analyses showed that better performance was associated with greater reliance on centroid-based gaze strategies, consistent with distributed monitoring of target configurations. Critically, older adults relied more on focal, target-based tracking under conditions simulating peripheral vision loss, and less on centroid-based strategies; this shift was associated with poorer performance. In contrast, oculomotor behavior during full-field viewing was largely preserved across age. Together, these findings suggest that aging affects multiple object tracking through combined sensory, attentional, and oculomotor mechanisms. Beyond a reduction in capacity, age-related decline also reflects systematic changes in visual sampling strategies during dynamic tracking.

## Introduction

Multiple object tracking (MOT) is a fundamental visual attention skill that enables observers to monitor several moving items simultaneously in dynamic environments and has been widely used to investigate the limits and mechanisms of dynamic visual attention in healthy observers (see e.g., Cavanagh & Alvarez, 2005 for a review). In young adults, successful tracking relies on the continuous attentional selection of several moving targets and is supported by efficient oculomotor strategies. Rather than maintaining fixation on a single target (e.g., Landry et al., 2001), successful tracking is often associated with smooth pursuit of the center of mass (or centroid) of the configuration formed by the targets combined with periodic saccadic recentering (Fehd & Seiffert, 2010; Fehd & Seiffert, 2008; Zelinsky & Neider, 2008, see also Hyönä et al., 2019 for a review). These strategies are thought to reduce spatial uncertainty and facilitate the maintenance of multiple object representations under dynamic attentional load.

A substantial body of work has demonstrated that MOT performance declines with advancing age, with older adults typically tracking fewer objects and exhibiting greater sensitivity to increases in task difficulty, such as those produced by higher object speeds or increased crowding (Sekuler et al., 2008; Trick et al., 2005). These impairments have generally been interpreted as reflecting age-related reductions in attentional capacity during dynamic visual tasks, particularly under conditions of increased attentional load (Sekuler et al., 2008). Aging has also been associated with reduced resistance to distraction and less efficient selective attention, factors that may further impair the ability to segregate targets from nearby distractors during tracking (Hasher & Zacks, 1988; Kramer et al., 1994). However, most existing studies on age-related changes in MOT have relied on cross-sectional comparisons between typically two discrete groups (younger vs. older adults), making it difficult to determine whether performance declines emerge gradually across the adult lifespan or instead reflect a more accelerated deterioration beginning at a particular age range. More recent work using broader age sampling suggests that age-related changes in motion and dynamic perception may follow nonlinear trajectories, with potential accelerations in later adulthood (Billino & Pilz, 2019). A finer-grained characterization of these trajectories is therefore needed to better understand how attentional tracking abilities evolve across adulthood.

In addition to attentional limitations, successful multiple object tracking (MOT) critically depends on the availability of visual information across the visual field. A large body of work in vision science has shown that central and peripheral vision provide complementary contributions to visual processing during natural viewing (Faurite et al., 2024; Larson & Loschky, 2009; Loschky et al., 2017; Nuthmann, 2014; Trouilloud et al., 2022). In particular, gaze-contingent paradigms simulating visual field loss, such as central scotoma (as in macular degeneration) or peripheral field restriction (as in retinitis pigmentosa), have demonstrated a functional dissociation between these two sources of visual input. Restricting central vision impairs fine spatial resolution and object identification, whereas restricting peripheral vision disrupts the extraction of global scene structure and the efficient guidance of eye movements during active vision (Findlay & Gilchrist, 2003; Land, 2006).

However, how alterations in visual field availability specifically affect MOT, and how these perceptual constraints may interact with age-related changes in tracking performance, remains largely underexplored.

Here, we investigated the effects of aging on MOT under both normal and altered viewing conditions. Sixty participants aged between 20 and 80 years completed a MOT task under full-field viewing and under gaze-contingent conditions selectively restricting either central or peripheral vision. To better characterize changes across the adult lifespan, we adopted a continuous sampling approach, recruiting exactly five participants per five-year age bin. Based on the literature reviewed above, we hypothesized that older adults would show reduced tracking performance across all viewing conditions. In addition, we expected age-related differences in tracking strategies, with older adults showing reduced efficiency in the use of grouping-based strategies during object tracking and a greater reliance on local or item-by-item tracking.

## Materials and methods

### Participants

Sixty participants aged 20 to 80 years (30 males, 30 females; 5 participants per five-year age group; see the cumulative age distributions in Figure 1) were involved in the study. None had any ocular disease. To assess their visual acuity, we used the Sloan letters of the Freiburg Visual Test monocularly, one eye after the other. In order to evaluate their cognitive functions, they all underwent the Mini-Mental State Examination (MMSE). They all had a score above 23. Finally, they reported how many hours they spent weekly playing video-games over the last 6 months. Participants were considered players if they spent at least 6 hours playing per week. The research was conducted at the Centre de Recherche et Cognition (Toulouse, France). Written informed consent was obtained, and the protocol was approved by the French Ethics Committee of Sud-ouest et outre-mer 2 (2020-A02441-38). Figure 1 shows the cumulative age distribution of all participants (in purple, see panel A), with the distributions for males and females shown in blue and red, respectively (see panel B).

**Figure 1:**
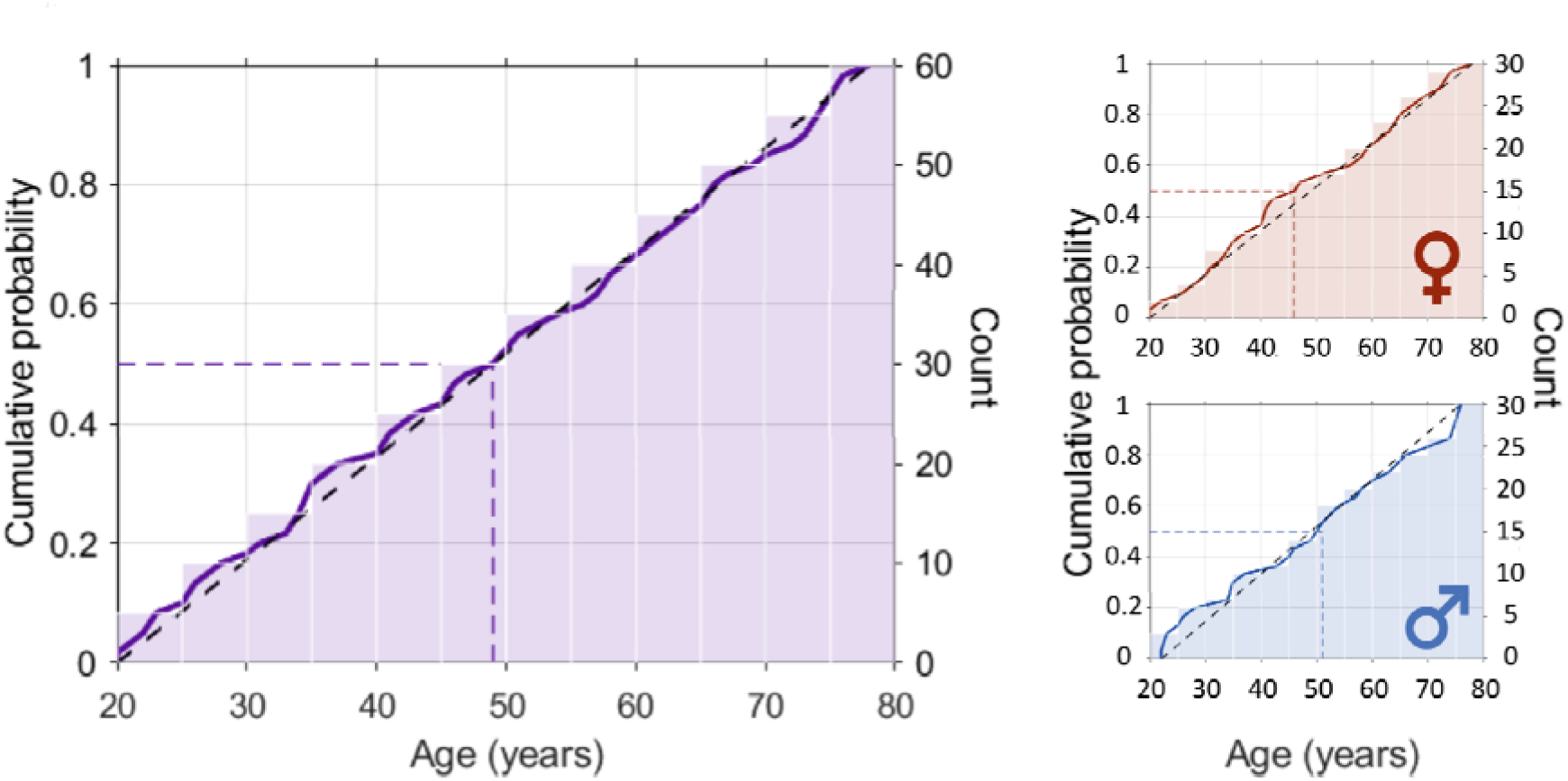
Cumulative age distribution of all participants (n=60, in purple) and corresponding distributions shown separately for females (red) and males (blue). Black dotted lines indicate the ideal case of a uniformly sampled population, whereas coloured dotted lines represent the medianage and cumulative probability values for each group.

### Experimental protocol

An overview of our experimental protocol is provided on Figure 2. Participants were seated in a chair with their head stabilized using a table-mounted support device equipped with chin and forehead rests (Figure 2-A). Visual stimuli were generated using Python (https://www.python.org/) and were projected onto a large convex screen (60Hz refresh rate, 1024x768 resolution) in a dark room at a viewing distance of 180 cm from the observers. The design of the stimuli was adapted from a previous study on multiple object tracking (Pylyshyn & Storm, 1988). The display consisted of six black disks (diameter: 2.8° of visual angle) moving on a white background. A frame of 6° of visual angle was added at the edges of the screen. The disks moved within an arena spanning 45° × 30° of visual angle. At the beginning of each trial, the six disks were randomly distributed on the screen without overlap. Three of them were randomly designated as targets and turned red, while the remaining three disks served as distractors (Figure 2-B). Participants were instructed to identify all target disks and then press a button to initiate the motion phase. The targets subsequently reverted to their original color, and all disks moved in random directions at a constant velocity of 7°/s for 10 s. This speed was chosen because it corresponds to the average preferred speed of neurons in the macaque middle temporal (MT) area, a region critically involved in motion processing (Priebe et al., 2006). When two disks collided, their trajectories were reassigned such that they moved in opposite directions along the line connecting their centers. When a disk collided with the boundary of the arena, its trajectory was reflected according to the angle of incidence. At the end of each trial, a single disk was highlighted in red while all other disks disappeared. Participants indicated via keyboard response whether the highlighted disk had been one of the three target disks (“a” key) or a non-target disk (“p” key), with chance performance at 50%. Auditory feedback signaled response accuracy. The protocol comprised five experimental conditions (Figure 2C). In the main condition (full-field), object tracking was performed under normal viewing. In the remaining conditions, a gaze-contingent paradigm was used to selectively mask either central or peripheral vision (see next section for details). In both cases, scotomas or apertures with radii of 10° and 15° were applied. Each participant completed 40 trials in each condition (full-field, 10° scotoma, 15° scotoma, 10° aperture, and 15° aperture), resulting in a total of 200 trials per participant. Condition order was randomized across participants. Participants performed the experiment monocularly with their best eye, while the other was patched. The entire experiment lasted approximately 90 minutes, and participants were free to take breaks at any time, as each trial was initiated only after a button press. In total, the dataset comprised 12,000 trials collected from 60 participants.

**Figure 2:**
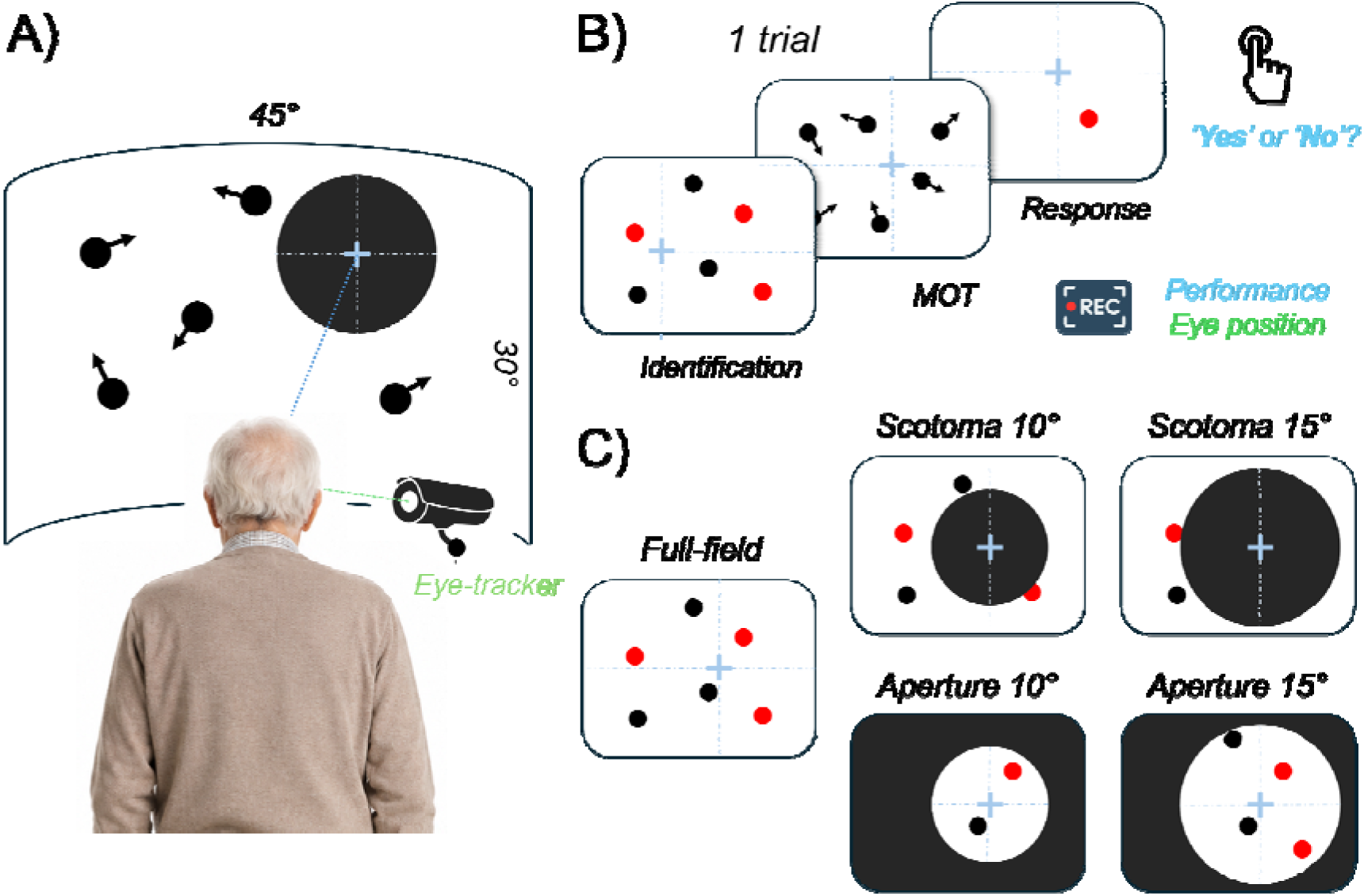
Protocol overview. **A) Experimental setup.** Participants tracked moving black disks projected onto a screen positioned 180 cm away while their eye movements were recorded. **B) Trial procedure.** At the start of each trial, the three target disks turned red and remained highlighted until the participant initiated the tracking phase by pressing a button. The targets then reverted to their original color, and all disks moved for 10 s, each being assigned a random initial direction of motion. At the end of the movement period, one disk was highlighted in red, and participants indicated whether it had been one of the targets. Each participant completed 40 trials in each of the five experimental conditions (200 trials in total), while both tracking performance and eye movements were recorded. **C) Experimental conditions.** The protocol comprised five distinct experimental conditions. The main condition was performed under normal viewing. In the remaining conditions, a gaze-contingent paradigm was used to selectively mask either central or peripheral vision. Scotomas and apertures with radii of 10° and 15° were applied in both cases.

### Gaze contingent paradigm

Eye movements were recorded monocularly using an EyeLink 1000 eye tracker (SR Research), sampled at 1000 Hz and positioned 30 cm in front of the participant. To simulate altered viewing conditions characteristic of clinical populations such as patients with macular degeneration or retinitis pigmentosa, a gaze-contingent paradigm was employed (see e.g., McIlreavy et al., 2012 or Janssen & Verghese, 2015). Central vision loss was simulated using a circular black disk with a radius of either 10° or 15° of visual angle, which was overlaid on the display and updated in real time to follow the participant’s gaze using the method proposed by Aguilar and Castet (2011). Similarly, central aperture conditions were simulated using a circular gaze-contingent aperture with a radius of either 10° or 15° of visual angle. In this condition, only the central portion of the visual field within the aperture remained visible, while the surrounding area was masked in black (Figure 2-C). We chose to use black masks to facilitate segmentation against the white background of the arena. A known limitation of gaze-contingent setups arises during blinks (Aguilar & Castet, 2011). As the upper eyelid begins to close, the eye tracker may register a downward displacement of the pupil centroid, resulting in a biased estimate of gaze position. Consequently, the artificial scotoma (or aperture) can be briefly displaced downward despite the absence of an actual eye movement. In addition, once the eyelid fully closes, the pupil is no longer detected and the gaze-contingent mask is typically removed. In principle, participants may exploit this brief period to view the display without masking. To mitigate these issues, blinks were identified as time periods during which pupil size fell below a predefined threshold (<100). During these intervals, the scotoma was no longer updated based on eye position and was instead maintained at the last valid gaze-contingent location. This procedure prevented spurious shifts of the mask during blinks and ensured that the scotoma remained stable and correctly positioned. To implement the gaze-contingent paradigm, each participant first completed a standard five-point calibration procedure, followed by a validation phase in which fixation accuracy was assessed across multiple targets. Calibration was accepted only when the mean spatial error was below 1° of visual angle.

### Oculomotor strategies

To characterize participants’ tracking strategies, each eye-position sample was classified according to the stimulus location closest to the gaze position based on Euclidean distance. Specifically, gaze samples were assigned to one of the three targets, to the targets’ centroid (i.e., center of mass), or to one of the distractors. Samples for which gaze position was more than 5° away from all of these locations were left unclassified. For each trial, the proportion of samples assigned to each category was calculated as a percentage of the total number of valid gaze samples. These analyses were conducted only for the full-field and aperture conditions, as they are not informative in the artificial scotoma conditions, where the center of gaze is masked and objects near fixation are therefore not visible. For the two artificial scotoma conditions, we instead quantified the proportion of time during which at least one of the three targets, or their centroid, was located within the visible portion of the visual field (i.e., outside the artificial scotoma).

We also conducted additional analyses of oculomotor strategies, quantifying the number of smooth pursuit eye movements (directed either toward one of the targets or toward their centroid), saccades, and gaze fixations, and examining whether these metrics were affected by age and/or conditions.

### Statistical analyses

All statistical analyses were conducted in RStudio (R version 4.3.1). The raw dataset comprised trial-level binary detection outcomes (detected vs. not detected) from the 12,000 valid trials completed by 60 participants across the five visual conditions.

Because the dependent variable consisted of binary responses and multiple observations were collected from each participant, detection performance was analyzed using binomial generalized linear mixed models (GLMMs) fitted with the *lme4* package (Bates et al., 2015). Age was mean-centered (Age_c) to provide an interpretable intercept and reduce multicollinearity in interaction terms, and was entered as a continuous predictor. Condition was entered as categorical predictors. Participant was included as a random-effect grouping factor using a random intercept.

Based on theoretical considerations, sex, visual acuity, video-game experience, and polynomial age terms were initially evaluated as potential covariates. Candidate models were compared using likelihood ratio tests (LRTs) and Akaike Information Criterion (AIC) values. Details of the model selection procedure are provided in Table 1.

**Table 1:** Model selection. Candidate GLMMs predicting detection performance were compared sequentially using likelihood ratio tests (LRTs). At each step, the current model was evaluated against the previously retained model, and was retained only if it provided a significantly better fit and a lower Akaike Information Criterion (AIC).

| Models | Fixed effects | AIC | LRT | p | Retained ? |
| --- | --- | --- | --- | --- | --- |
| M1 | Age, Condition | 14086 | - | - | Yes |
| M2 | Age * Condition | 14050 | 43.5 | < 0.001 | Yes |
| **M3** | **M2 + Sex** | **14045** | **7** | **0.008** | **Yes** |
| M4 | M3 + Acuity | 14049 | 0 | - | No |
| M5 | M3 + VideoGame | 14046 | 1.4 | 0.24 | No |
| M6 | M3 + Order | 14046 | 1.6 | 0.202 | No |
| M7 | (Age_c + Age_c<sup>2</sup>) * Condition + Sex | 14053 | 2 | 0.854 | No |

The Age_c × Condition interaction and Sex significantly improved model fit and were therefore retained. Visual acuity, video-game experience, and polynomial age terms did not improve model fit and were therefore excluded from the final model. In the final model, Full-field and Female were used as reference levels for Condition and Sex, respectively. The primary analysis of detection performance was based on the following model:

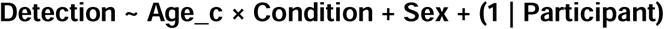

To further characterize Age_c × Condition interactions, age-related slopes were estimated separately for each viewing condition using the *emtrends* function from the *emmeans* package. Planned contrasts were then performed to directly compare age-related slopes between conditions.

A second analysis aimed at evaluating the influence of different gaze strategies on the detection performance. To avoid introducing noise into the statistical models, we excluded trials containing fewer than 75% valid gaze-position samples. Of the 12,000 recorded trials, 10,983 trials were retained, corresponding to 91.5% of the original dataset. To assess the impact of this threshold choice on the results, all analyses were repeated using alternative thresholds of 65% and 85%. The overall pattern of results remained unchanged across thresholds. Because the eye tracker sampling rate (1000 Hz) differed from the screen refresh rate (60 Hz), only eye-tracking samples temporally aligned with the display refresh frequency were retained for analysis. Because gaze position relative to targets and centroids is not informative in the artificial scotoma conditions, analyses involving gaze-allocation measures were restricted to a reduced dataset including only the Full-field, Aperture10, and Aperture15 conditions. Oculomotor strategy measures (i.e., the percentage of time spent looking at targets or the centroid) were z-standardized within the analyzed dataset, yielding the variables Target_z and Centroid_z. It should be noted that Target_z and Centroid_z were not negatively correlated. Although target-looking and centroid-looking may appear to represent opposite oculomotor strategies, participants could alternate between these strategies within the same trial rather than relying exclusively on one of them. Consequently, the two variables captured partially independent aspects of gaze behavior and could be entered simultaneously in the model without problematic multicollinearity. Detection performance was reanalyzed on this reduced dataset (N = 6,840 trials) using a new GLMM structure, defined as follows:

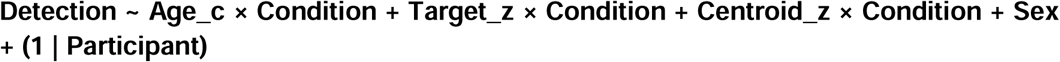

Finally, to determine whether gaze-allocation strategies varied with age and viewing condition, and whether these changes could account for age-related differences in tracking performance, separate models were fitted with Target_z and Centroid_z as dependent variables. Because these variables consisted of continuous z-standardized proportions measured repeatedly across trials within participants, linear mixed-effects models (LMMs) fitted with the *lmerTest* package were used. The models were specified as follows:

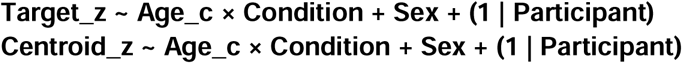

## Results

In this study, we investigated the effects of aging on multiple object tracking (MOT) under both normal and altered viewing conditions. A gaze-contingent paradigm was used to simulate visual impairments characteristic of clinical populations, such as central vision loss in macular degeneration and peripheral vision loss in retinitis pigmentosa. Performance and eye movements were recorded in a sample of 60 participants aged between 20 and 80 years. In the following, we first describe the effects of aging on behavioural performance before examining its impact on oculomotor strategies.

### Effects of age and visual condition on detection performance

Figure 3 shows the percentage of correct tracking for each participant as a function of age. Results are first presented across all conditions (upper-left panel), followed by the full-field condition (blue, bottom-left panel), the two artificial scotoma conditions (orange, middle panels), and the central aperture conditions (purple, right panels). The GLMM revealed a significant main effect of centered age (β = −0.028, SE = 0.004, *p* < 0.001), indicating that performance decreased with increasing age. The model also revealed a significant main effect of condition (*p* < 0.001). Relative to Full-field condition (reference level in the GLMM), detection performance was lower in the Small Scotoma (β = −1.899, SE = 0.08, p < .001), Large Scotoma (β = −1.959, SE = 0.08, p < .001), Large Aperture (β = −1.233, SE = 0.08, p < .001), and Small Aperture (β = −1.610, SE = 0.08, p < .001) conditions. Post hoc comparisons further revealed that detection performance was significantly lower in the two scotoma conditions than in either aperture condition (all *p*_holm < .001), whereas the two scotoma conditions did not differ from each other (*p*_holm = .30). The two aperture conditions also differed significantly from each other (*p*_holm < .001), with lower performance in the Small than in the Large Aperture condition. Mean detection probabilities estimated from the final model (marginalized over participant age and sex, with 95% confidence intervals) were as follows: Full-field condition (M = 0.900, 95% CI [0.886 : 0.913]), followed by the Large Aperture (M = 0.724, [0.701 : 0.746]), Small Aperture (M = 0.643, [0.618, 0.667]), Small Scotoma (M = 0.575, [0.549 : 0.600]), and Large Scotoma (M = 0.560, [0.534 : 0.585]). Finally, a significant Age × Condition interaction was observed. Follow-up analyses indicated that this interaction was driven by differences in the effect of Age between the Full-field condition and Small Scotoma (β = 0.023, SE = 0.005, *p* < 0.001), Large Scotoma (β = 0.024, SE = 0.005, *p* < 0.001), and Small Aperture (β = 0.020, SE = 0.005, *p* = 0.001) conditions. In contrast, the difference in the age effect between the Full-field and Large Aperture conditions was not significant (β = 0.009, SE = 0.005, *p* = 0.075). This interaction pattern may reflect floor effects, as performance in the masked conditions approached chance level and therefore could not decrease substantially further. To directly assess whether age-related declines differed between central and peripheral visual restrictions, we compared the age slopes observed in the Aperture and Scotoma conditions. Contrasting the two aperture conditions with the two scotoma conditions revealed a steeper age-related decline in the apertures (β = 0.009, SE = 0.002, p < 0.001), consistent with the idea that performance in the scotoma conditions is constrained by a floor effect that masks age-related differences. To address this, we compared conditions individually. The contrast between the Large Aperture and the Small Scotoma, the two least floor-bound conditions, confirmed a significantly steeper age-related decline in the aperture (β = 0.014, SE = 0.004, p < 0.001). In contrast, the comparison between the Small Aperture and the Large Scotoma, both closer to floor-bound conditions, was not significant (β = 0.004, SE = 0.003, p = 0.28). Together, these results suggest that the age-related decline is more pronounced when central rather than peripheral vision is restricted, but that this difference is partially obscured when both conditions approach floor performance.

**Figure 3:**
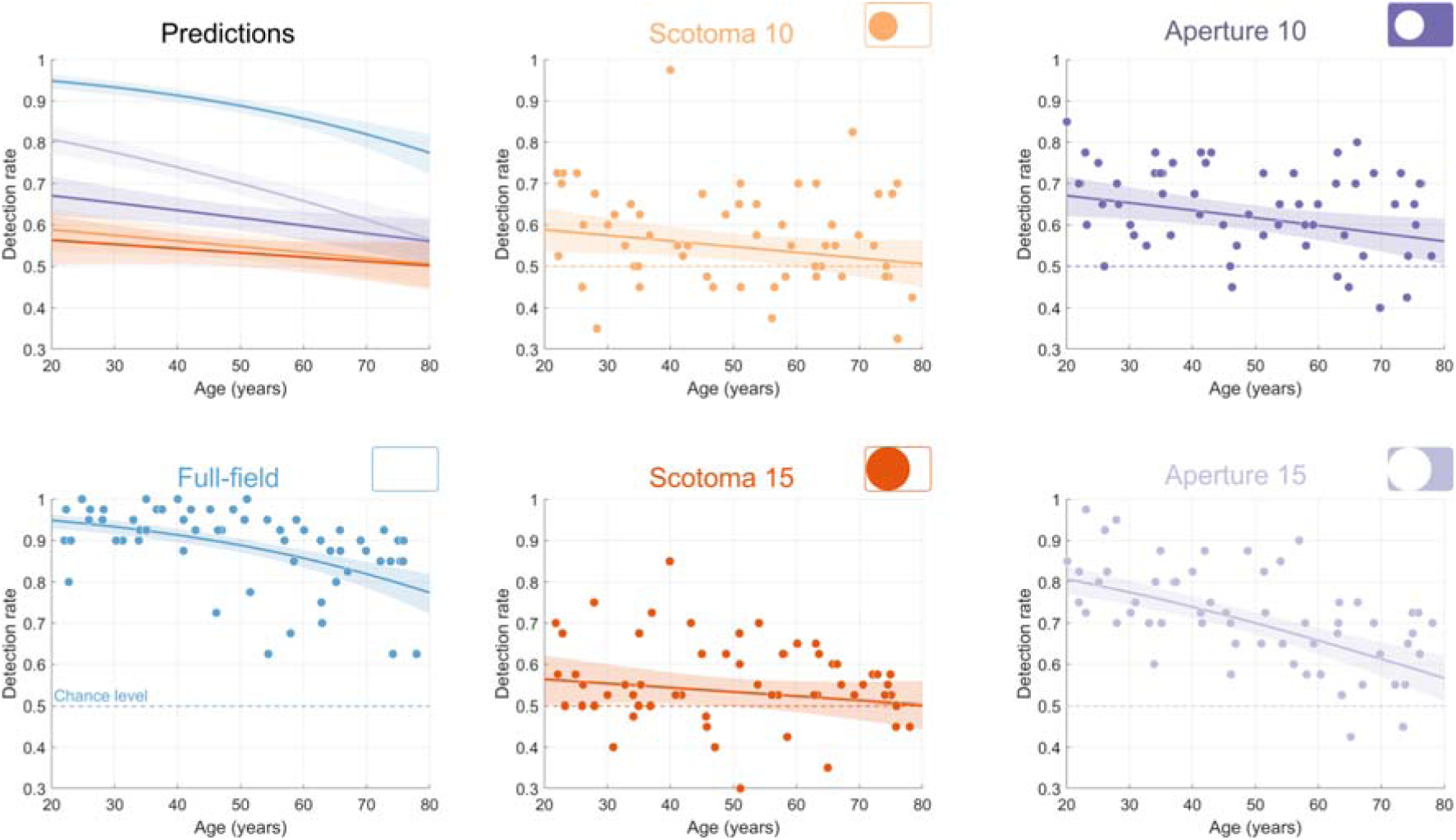
Percentage of correct tracking as a function of age. The upper-left panel presents model predictions for all conditions simultaneously (Full-field in blue; 10° and 15° artificial scotomas in light and dark orange, respectively and 10° and 15° apertures in dark and light purple, respectively). Colored lines indicate the estimated relationship between performance and age, and shaded regions represent the corresponding 95% confidence intervals. Results are also shown separately for each condition. In these panels, each point represents an individual participant’s observed performance averaged across 40 trials. For visualization purposes, a small horizontal jitter was applied to reduce point overlap.

Taken together, these findings suggest that aging impairs visual tracking performance and that restrictions of the visual field further reduce performance. However, the effect of age under masked conditions appears to be attenuated, likely due to floor effects arising from increased task difficulty.

### Effects of oculomotor strategy

To determine whether fixation strategies were associated with detection performance independently of age, we examined whether the proportion of time spent fixating either the targets themselves (*Target_z*) or their centroid (*Centroid_z*; i.e., center of mass) predicted trial outcome. Fixating the centroid is generally thought to reflect a distributed attentional strategy that sacrifices precise foveal encoding of individual targets in favor of more stable, global monitoring of the target set. The relationship between these z-scores and performance is shown in Figure 4A (left panel: Target tracking; right panel: Centroid tracking). The GLMM revealed a significant positive effect of Centroid_z on detection performance (β = 0.353, SE = 0.087, p < .001), indicating that greater fixation of the centroid was associated with better performance in the Full-field condition. In contrast, Target_z was not significantly associated with better performance in the Full-field condition (β = 0.111, SE = 0.102, p = .27). There were no significant interactions between Target_z and either central aperture conditions (Small Aperture: β = 0.175, SE = 0.118, p = .138; Large Aperture: β = −0.020, SE = 0.125, p = .875), nor between Centroid_z and either central aperture conditions (Small Aperture: β = −0.194, SE = 0.112, p = .085; Large Aperture: β = −0.110, SE = 0.108, p = .310).

**Figure 4:**
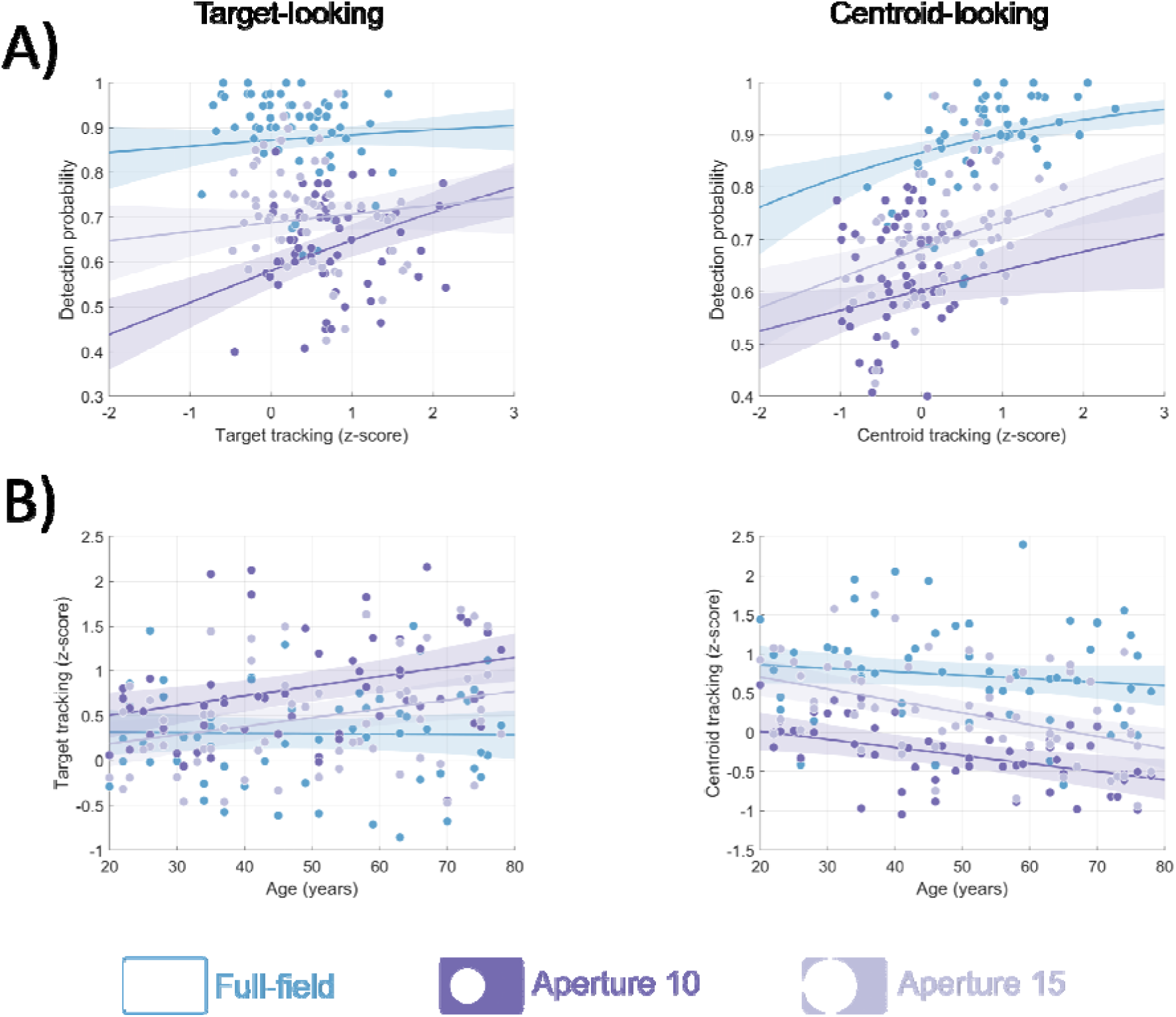
Association with oculomotor strategy. (A) Percentage of correct responses plotted as a function of the time spent looking at any of the three targets (left panel) or at their centroid (right panel). Viewing times were standardized and expressed as Z-scores. Results are shown for the full-field condition (blue) and the two central aperture conditions (purple). Each dot represents the mean performance of an individual participant across valid trials. Colored lines indicate the estimated relationship between performance and Z-scores. (B) Tracking strategy (Z-scores) as a function of age.

There was no significant interaction between Target_z or Centroid_z (all *p* > 0.08) and either aperture conditions, suggesting that the effects observed in the Full-field condition were consistent across conditions. Nevertheless, inspection of condition-specific slopes (computed via emtrends) provides a complementary view as target looking emerged as a significant positive predictor in the Small Aperture condition (β = 0.287, SE = 0.060, 95% CI [0.170 : 0.405]). The pattern in the Large Aperture was intermediate and non-significant (β = 0.092, SE = 0.074, 95% CI [−0.052 : 0.236]). Together, these findings are consistent with the hypothesis that the optimal oculomotor strategy depends on the available visual field. The centroid strategy, classical in multiple object tracking (Fehd & Seiffert, 2008), remains effective across the three conditions; In addition, when the visual field is severely restricted (small aperture), target-following becomes an effective complementary strategy.

To determine whether oculomotor strategies varied with age in the Full-field and Aperture conditions, and whether they could account for the poorer performance observed in older participants, we fitted two LMMs, one for each strategy (centroid- and target-looking; see Figure 4-B). In the full-field condition, age had no significant effect on either strategy (centroid: β = −0.004, SE = 0.003, 95% CI [−0.011 : 0.002], p = 0.19; target: β = −0.0004, SE = 0.003, 95% CI [−0.007 : 0.006], p = 0.90), indicating that younger and older participants allocated comparable amounts of time to each gaze pattern when full vision was available. In contrast, significant effects of age on both strategies were found in the aperture conditions (centroid, small aperture: β = −0.006, SE = 0.001, p < 0.001; large aperture: β = −0.011, SE = 0.001, p < 0.001; target, small aperture: β = +0.011, SE = 0.001, p < 0.001; large aperture: β = +0.010, SE = 0.001, p < 0.001). To interpret these effects directly, condition-specific age slopes were computed (*emtrends*). With increasing age, time spent on the centroid declined progressively under restricted vision (small aperture: β = −0.010, SE = 0.003, 95% CI [−0.017 : −0.004]; large aperture: β = −0.015, SE = 0.003, 95% CI [−0.022 : −0.009]), while time spent on individual targets increased (small aperture: β = +0.011, SE = 0.003, 95% CI [0.004 : 0.018]; large aperture: β = +0.010, SE = 0.003, 95% CI [0.003 : 0.017]). This pattern suggests that, under restricted viewing conditions, older participants progressively shift from a grouping (centroid-based) strategy, which our first analysis identified as the more efficient strategy in these conditions, towards a more focal, target-by-target strategy, and that this shift may contribute to their poorer performance.

Several studies on multiple object tracking have suggested that the most effective oculomotor strategy may involve tracking a weighted centroid, referred to as the anticrowding point, that shifts toward targets nearest to distractors, thereby reducing the effects of crowding (Lukavský, 2013). To assess whether this strategy improved performance and whether its use differed across age groups, we replicated the statistical analyses described above. Detected gaze positions were classified as falling on a target, the target centroid, or the anticrowding point. Because these regions could overlap, each sample was assigned to the nearest point of interest. As the proportion of time spent on the centroid and on the anticrowding point was highly correlated, including both predictors in the same model would have introduced multicollinearity and reduced model reliability. We therefore fitted an alternative model with the same structure, replacing centroid time with time spent on the anticrowding point. However, given the strong correlation between these variables, results did not differ from those reported in the previous paragraph.

As noted above, all these analyses of oculomotor strategies are not informative in the artificial scotoma conditions, where the center of gaze is masked and objects near fixation are therefore not visible. For these conditions, we instead quantified the proportion of time during which at least one of the three targets, or their centroid, was located within the visible portion of the visual field (i.e., outside the artificial scotoma). In this case, a new GLMM including z-standardized visibility indices as predictors, together with their interactions with condition, revealed no significant effect of target or centroid visibility on performance. However, increasing age was associated with reduced target and centroid visibility, particularly under the most restricted viewing condition. Given the absence of a significant relationship between visibility measures and performance, all associated analyses are reported in the Supplementary Materials (see ***Supplementary materials 1***).

### Gender differences

Sex was retained in the final model because its inclusion significantly improved model fit. Male participants showed higher odds of providing a correct response than female participants (β = 0.21, SE = 0.08, p = 0.006), corresponding to a 24% increase in odds for a given age (OR = 1.24). To determine whether this effect reflected a general performance difference or whether age and visual condition influenced men and women differently, we fitted an additional GLMM including interactions between Sex, Age, and Condition. No significant interactions were observed, and this extended model did not improve model fit relative to the previous model (χ²(5) = 5.99, *p* = 0.31).

### Additional analyses

As described in the Methods section, we further analyzed participants’ eye movements by quantifying the number of smooth pursuit movements (directed either toward one of the targets or toward their centroid), saccades, and gaze fixations. Because these analyses did not yield significant effects on performance, the associated methodology and results are provided in the supplementary materials (see ***Supplementary Material 2***).

Finally, to assess the robustness of our experimental design, we examined whether presentation order influenced performance. No substantial order effects were observed (see ***Supplementary Material 3***). However, a modest learning effect emerged in the central aperture conditions: participants who were exposed to the wider aperture before the narrower one showed improved performance in the more demanding condition.

## Discussion

The present study investigated how healthy aging affects multiple object tracking (MOT) under normal viewing conditions and under gaze-contingent manipulations simulating central and peripheral visual field loss. By combining a lifespan sampling strategy (see Figure 1) with eye-tracking measures of oculomotor behavior (Figure 2), we sought to characterize not only whether tracking performance declines with age, but also how aging influences the visual strategies used to support dynamic attentional tracking. Three main findings emerged. First, tracking performance declined progressively across adulthood (Figure 3), confirming that MOT is highly sensitive to age-related changes in visual-attentional processing. Second, restricting visual-field availability strongly impaired performance, with artificial central scotomas producing the largest behavioral costs. Third, age-related performance declines were accompanied by changes in oculomotor strategy under restricted viewing conditions (Figure 4), with older participants relying more heavily on focal target-based tracking and less on centroid- or grouping-based strategies associated with better performance.

### Aging and dynamic attention tracking

Our results are consistent with previous studies showing age-related impairments in MOT and dynamic visual attention (Sekuler et al., 2008; Trick et al., 2005). Importantly, the present study extends this literature by characterizing these changes continuously across the adult lifespan rather than through categorical comparisons between younger and older adults. The progressive decline observed here suggests that MOT abilities deteriorate gradually with age, consistent with broader evidence showing continuous age-related changes in motion and dynamic perception (Billino & Pilz, 2019). Several mechanisms may contribute to this decline. MOT requires sustained allocation of attention across multiple moving objects while resisting interference from nearby distractors. Aging has been associated with reduced attentional capacity, slower processing speed, and impaired inhibitory control (Hasher & Zacks, 1988; Kramer et al., 1994), all of which may compromise the maintenance of stable target representations in crowded dynamic displays. In addition, reduced motion sensitivity and temporal processing may impair the efficient updating of target positions over time.

Age-related changes in low-level vision may also contribute to poorer MOT performance. Visual acuity declines with age, and our sensitivity analyses confirmed that acuity predicts detection performance under unrestricted viewing conditions; however, this effect is abolished in the presence of a central scotoma. Because age and acuity were strongly correlated in our sample, including both predictors did not improve model fit, and their respective contributions could not be formally dissociated. Nevertheless, age-related impairments were accompanied by systematic changes in oculomotor strategy under restricted viewing conditions, which cannot be attributed solely to sensory decline, suggesting that the observed decline in performance reflects an interaction between sensory, attentional, and oculomotor factors.

As reported in the result section, the effect of age was attenuated in the most difficult gaze-contingent conditions, particularly those involving artificial scotomas or the smallest apertures. These attenuations likely reflect a floor effect rather than preserved processing as performance approached chance level and could not decrease further.

### Contributions of central and peripheral vision to multiple object tracking

The gaze-contingent manipulations revealed that MOT depends strongly on both central and peripheral visual information. The largest deficits occurred in the artificial scotoma conditions, suggesting a particularly important role for central vision. While previous work has emphasized peripheral vision’s contribution to scene perception and eye-movement guidance (Findlay & Gilchrist, 2003; Land, 2006), MOT also requires the continuous updating of target positions and the resolution of close target-distractor interactions.The strong impact of central masking suggests that central vision supports both the maintenance of stable target representations and the resolution of spatial interference, contributing more broadly to attentional allocation and the online monitoring of dynamic target-distractor relationships. Given the large size and high saliency of the stimuli, reduced visibility alone is unlikely to account for this effect. Central aperture conditions also substantially impaired performance, especially with the smallest aperture, suggesting that peripheral vision contributes to the global monitoring of target configurations and the anticipation of interactions outside the foveal region. Peripheral information may therefore support the distributed attentional processes required to maintain a coherent representation of the target group. Together, these results indicate that MOT depends on complementary interactions between central and peripheral visual processing rather than on either source alone.

These findings may also help explain difficulties experienced by clinical populations with visual field defects. Patients with macular degeneration often report difficulties tracking moving objects despite preserved peripheral vision, whereas patients with retinitis pigmentosa show impairments in navigation and dynamic scene monitoring because of peripheral field restriction. Although simulated scotomas cannot reproduce the long-term adaptations observed in these patients, the present results suggest that both forms of visual field loss substantially disrupt the attentional mechanisms supporting MOT.

### Oculomotor strategies and aging

A central objective of the study was to determine whether age-related reductions in tracking performance were associated with changes in oculomotor strategy. Consistent with previous eye-tracking studies of MOT (Fehd & Seiffert, 2008, 2010), participants who spent more time fixating the centroid of the target configuration showed better tracking performance (Figure 4), supporting centroid-looking constitutes as an efficient strategy for monitoring multiple moving targets simultaneously. Maintaining gaze near the geometric center of the target group may reduce spatial uncertainty, minimize the need for large saccades, and maintain all targets within a relatively homogeneous region of visual sensitivity. Importantly, this strategy appeared to be altered with aging under restricted viewing conditions. Whereas younger participants continued to allocate substantial gaze time to the centroid, older adults increasingly fixated individual targets directly, particularly in the central aperture conditions. This shift toward focal target tracking was accompanied by reduced performance, suggesting that older adults relied less effectively on distributed grouping-based strategies. One possible interpretation is that aging reduces the efficiency with which observers can maintain integrated representations of multiple targets. Grouping-based strategies likely depend on the ability to encode spatial relationships among objects and dynamically update these relationships over time. Age-related reductions in attentional flexibility or visuospatial working memory may therefore bias older observers toward more local item-by-item monitoring strategies (Verhaeghen & Cerella, 2002). Although such strategies may provide more precise information about individual targets, they are likely less efficient when multiple objects must be monitored simultaneously in dynamic environments. This account is consistent with evidence showing age-related declines in visual-attentional efficiency under conditions requiring information to be processed across multiple spatial locations (Seiple et al., 1996; A. B. Sekuler et al., 2000).

### Sex difference in MOT

Although sex differences were not the primary focus, male participants showed modestly higher tracking performance than female participants across conditions, broadly consistent with previous psychophysical studies reporting sex-related differences in motion processing and visuospatial tasks (e.g., Jin et al., 2023; Murray et al., 2018; Shaqiri et al., 2018 or Guénot et al., 2023). Importantly, however, no interactions involving sex and age or viewing conditions were observed, suggesting that the mechanisms underlying the effects of aging and visual-field restriction were qualitatively similar in males and females. The origin of sex differences in motion-related tasks remains debated. Potential contributors include individual differences in visuospatial abilities, attentional allocation, action-oriented experience, and video-game practice. Gaming experience did not explain additional variance and was excluded from the final model, partly because gamers were few and predominantly male (9/11). Future studies specifically targeting individual differences in MOT performance could clarify the respective roles of perceptual, cognitive, and experiential factors.

### Methodological considerations and limitations

Several limitations should be acknowledged. The task involved tracking only three targets among six objects at a single speed. Manipulating target load or motion speed could reveal additional interactions between aging and visual-field restrictions, as older adults may show disproportionate impairments under higher attentional demands (Bettencourt & Somers, 2009; Pylyshyn & Storm, 1988; Trick et al., 2005; Yantis, 1992) or faster velocities (Alvarez & Franconeri, 2007; Franconeri et al., 2008). In addition, performance estimates were based on a relatively limited number of trials (40 per condition), a choice driven by the need to limit overall experiment duration and participant fatigue, particularly given the wide age range and the demands of the gaze-contingent paradigm. While sufficient to reveal robust effects of age and condition, a greater number of trials would improve the reliability of individual estimates and support fine-grained analyses of variability.

Although gaze-contingent paradigms provide a powerful means of simulating visual field loss, they cannot fully replicate the perceptual, behavioral, and neural adaptations associated with chronic ophthalmologic disease. Individuals with long-standing central or peripheral visual impairments often develop compensatory mechanisms, such as preferred retinal loci (see, e.g., Tarita-Nistor et al., 2023) and modified visual scanning strategies (Verghese et al., 2021; Wiecek et al., 2012), which may be absent in healthy observers exposed to simulated deficits. Nevertheless, a recent study using a similar experimental paradigm reported comparable tracking performance in a small sample of four participants with macular degeneration (MD) and age-matched control participants tested with a gaze-contingent artificial scotoma (Michaud et al., 2026), suggesting that gaze-contingent paradigms can provide a useful approximation of the functional impact of central visual field loss on MOT performance.

## Conclusion

We found that multiple object tracking declines progressively across the adult lifespan and is strongly modulated by the availability of central and peripheral visual information. Beyond reduced performance, aging is associated with systematic changes in oculomotor strategies under visually constrained conditions simulating peripheral vision loss, with older adults relying less on efficient centroid-based grouping strategies. These findings highlight the interplay between sensory constraints, attentional capacity, and adaptive changes in visual sampling in age-related tracking impairments.

## Ethics Statement

All participants gave their written informed consent and the protocol was approved by the French Ethics Committee (CPP) of Sud-Ouest et Outre-Mer 2 (2020-A02441-38).

## Acknowledgements

We thank Clément Libreau for his help in the recordings.

## Supplementary materials

### 1- Supplementary analyses of oculomotor strategies in central scotoma conditions

For the two central scotoma conditions, we quantified the proportion of time during which at least one of the three targets, or their centroid, was located within the visible portion of the visual field (i.e., outside the artificial scotoma). We then fitted a GLMM including these two z-standardized visibility indices as predictors, together with their interactions with condition. Neither target visibility nor centroid visibility significantly predicted performance. Specifically, no effect of target visibility was observed in the small scotoma condition (β = 0.071, SE = 0.156, p = .66), and its interaction with the large scotoma condition was not significant (β = −0.054, SE = 0.160, p = .73). Likewise, centroid visibility showed no significant effect in the small scotoma condition (β = 0.033, SE = 0.059, p = .57) and no significant interaction with the large scotoma condition (β = −0.131, SE = 0.076, p = .09). These findings suggest that visibility measures did not account for performance differences across these conditions.

Next, to assess age-related changes in visibility, we fitted two LMMs with target visibility and centroid visibility as dependent variables. Age was not associated with target visibility in the small scotoma condition (β = −0.001, SE = 0.002, *p* = .68), but a significant interaction with the large scotoma condition emerged (β = −0.006, SE = 0.002, *p* < .001), showing that target visibility decreased with age under the large scotoma condition. Centroid visibility also decreased significantly with age in the small scotoma condition (β = −0.007, SE = 0.003, *p* = .043), with a further reduction in the large scotoma condition (interaction: β = −0.003, SE = 0.001, *p* = .04). Taken together, these results indicate that older participants spent less time with the targets and their centroid visible, especially under the most restricted viewing condition.

**Supplementary Figure 1.**
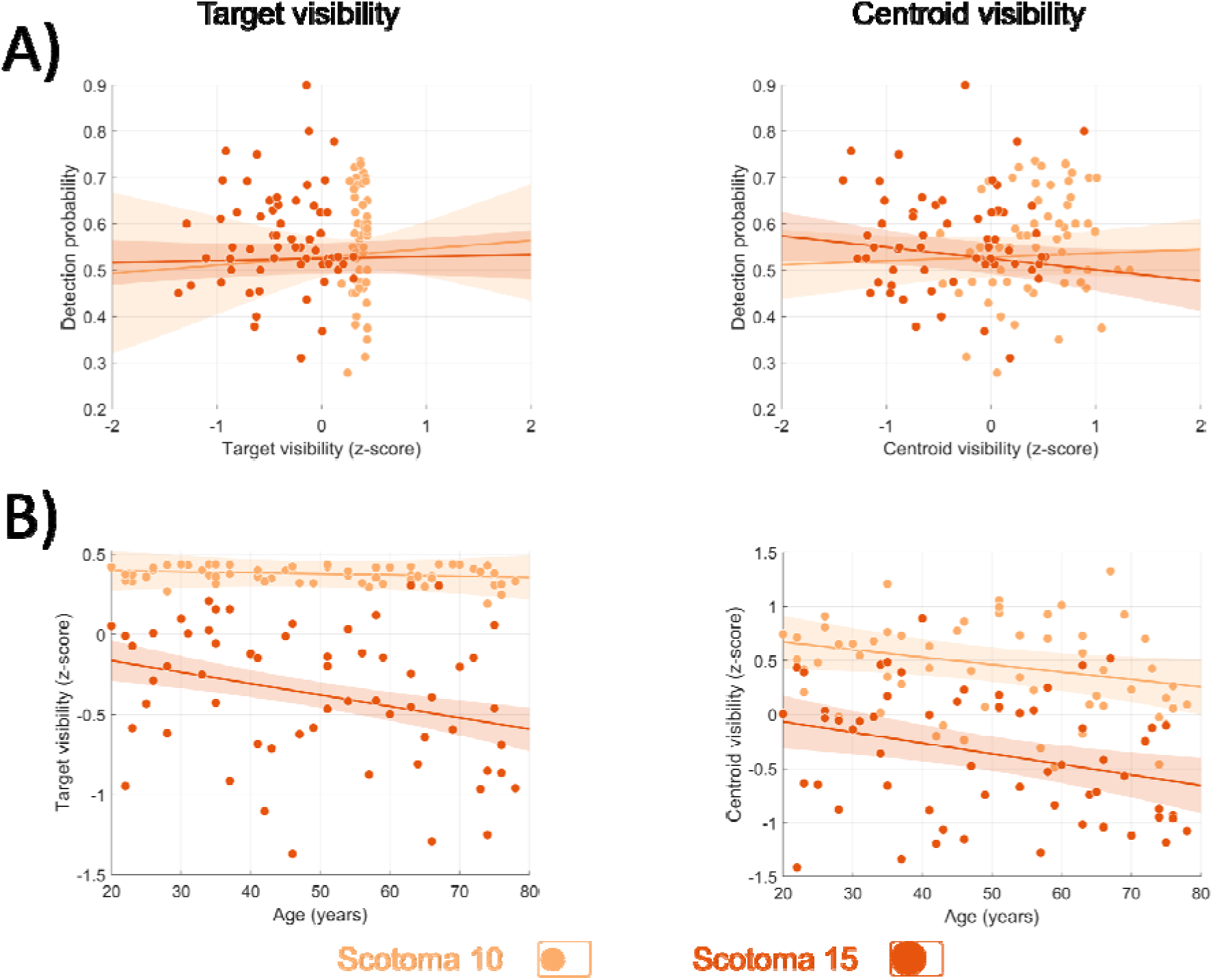
: Target and centroid visibility under artificial scotoma conditions. (A) Percentage of correct responses as a function of the proportion of time during which at least one target (left panel) or the centroid (right panel) was visible (i.e., located outside the scotoma). Visibility measures were z-standardized and expressed as Z-scores. Data are shown separately for the two scotoma conditions (light and dark orange). Each dot represents the mean performance of an individual participant across valid trials. Colored lines depict the model-estimated relationship between performance and visibility. (B) Tracking strategy (Z-scores) as a function of age.

### 2- Supplementary analyses of oculomotor strategies: Gaze fixation, saccades and smooth pursuit

To complete our analyses of oculomotor strategies, detected gaze-position signals were classified into three categories: saccades, smooth pursuit, and fixations. To identify fixations, raw gaze samples were first smoothed using a second-order zero-phase Butterworth low-pass filter with a cutoff frequency of 3 Hz, reducing the likelihood that small noise-related velocity fluctuations would be misclassified as fixations. Fixations were then defined as periods during which eye velocity remained below 1°/s. For saccade detection, the unfiltered raw eye-velocity signal was used directly. Saccades were identified using a velocity threshold of 30°/s combined with a displacement criterion of at least 1° of visual angle, following Cornelissen (2005). Smooth pursuit was identified from gaze samples with velocities between 5 and 9°/s when tracking targets, and between 1 and 3°/s when tracking the centroid. A new GLMM with eye movements (i.e. the percentage of time spent doing saccades, smooth pursuit or fixation) as predictors was fitted. These variables were z-standardized within the analyzed dataset, yielding the variables *Fixation_z*, Saccades_z, Pursuit_z and Pursuit_centroid_z with a mean of 0 and a standard deviation of 1. None of the eye movements significantly contributed to detection performance (all *p* >0.15) suggesting that the proportion of time spent fixating, doing smooth pursuit, or making saccades did not influence detection accuracy.

Next, we examined whether eye movements varied as a function of age. To this end, we fitted four linear mixed-effects models (LMMs) using the same structure as in the previous analyses, with each eye movement measure entered as the dependent variable. Regarding fixations, no main effect of age was observed in the full-field condition (*p* = 0.41). However, a significant interaction emerged between centered age and the large scotoma condition (β = 0.009, SE = 0.001, *p* < .001), as well as in the narrow central aperture condition (β = 0.004, SE = 0.001, *p* < .001), suggesting that older participants tended to make more fixations in these conditions. Regarding smooth pursuit of targets, no effect of age was observed in the full-field condition (*p* = 0.265). However, older participants showed significantly more pursuit movements in both central aperture conditions (narrow aperture: β = 0.006, SE = 0.0007, *p* < .001; large aperture: β = 0.005, SE = 0.0007, *p* < .001) and slightly fewer in the large scotoma condition (β = -0.002, SE = 0.0007, *p* = 0.018). Overall, these results suggest that older participants relied more on smooth pursuit in central aperture conditions specifically. Regarding smooth pursuit of the centroid, a significant main effect of age was observed in the full-field condition (β = -0.009, SE = 0.003, *p* < 0.001), indicating that older participants spent less time pursuing the centroid overall. This age-related reduction was amplified under scotoma conditions (narrow aperture: β = 0.006, SE = 0.001, *p* < 0.001; large aperture: β = 0.01, SE = 0.001, *p* < 0.001) but attenuated under the large central aperture condition (β = -0.003, SE = 0.0009, *p* = 0.003). Regarding saccades, no main effect of age was observed (*p* = 0.6). However, older participants made significantly fewer saccades across all restricted viewing conditions (all *p* < .001), with the strongest reduction observed under the narrow central aperture condition (β = -0.023, SE = 0.001, *p* < .001).

**Supplementary Figure 2:**
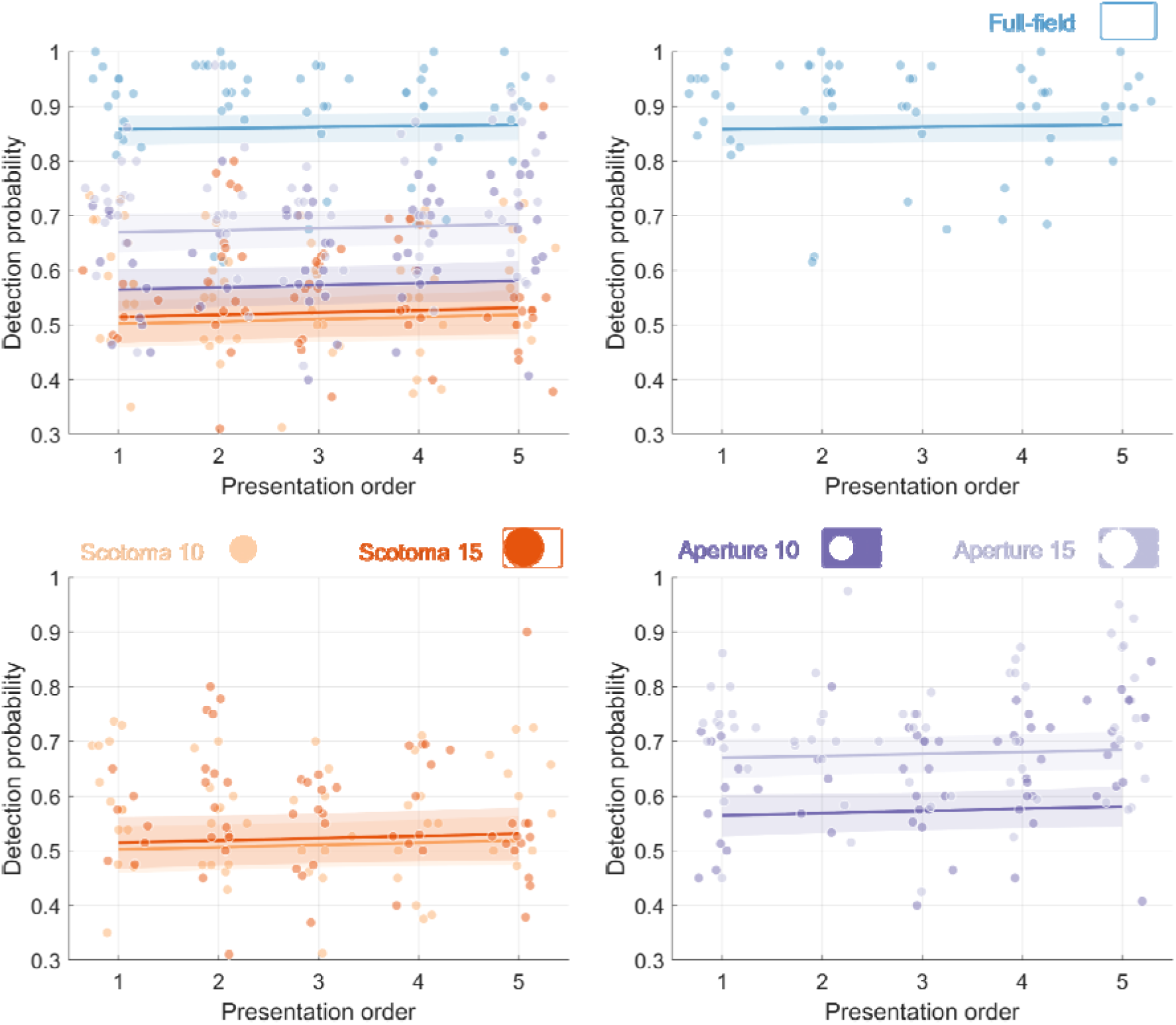
Effect of condition order on performance. The top left plot combines all three subplots where colored lines depict the model-estimated relationship between performance and presentation order. Each dot represents the mean performance of an individual participant across valid trials.

### 3- Supplementary analyses of conditions order

Because the order of conditions was randomized across participants, we examined whether it influenced performance. To this end, we fitted an additional model including condition order as a predictor. This analysis revealed no significant effect of order (β = 0.017, SE = 0.016, *p* = 0.29).

